# Antimicrobial activity of diverse chemotypes of *Lippia graveolens* against *Aeromonas hydrophila* isolated from tilapia

**DOI:** 10.1101/2023.07.24.549066

**Authors:** Josué García-Pérez, Juan Pérez-Sabino, Susana Mendoza-Elvira, Antonio Jorge Ribeiro da Silva, Juan Ulloa-Rojas

## Abstract

**Objetive:** This study aimed to evaluate the antimicrobial efficacy of essential oil (EO) from diverse chemotypes of *Lippia graveolens* against oxytetracycline-resistant *Aeromonas hydrophila*, which primarily affects the tilapia aquaculture (*Oreochromis* sp) in Guatemala.

**Methodology:** *L. graveolens* were collected in three departments in Guatemala, the EO was obtained by hydrodistillation and characterized by gas chromatography and mass spectrometry (GC/MS). Subsequently, an antimicrobial assay was conducted by screening the disk and dilution susceptibility tests, and evaluation of synergistic interactions among the chemotypes, each test being carried out in triplicate.

**Results:** The analysis revealed the presence of twenty-seven compounds in the EO obtained from the chemotypes, the main class being monoterpene. The major constituents identified were cis-Dihydro-β-terpineol (8.84%) in chemotype I, carvacrol (51.82%) in chemotype II, and thymol (79.62%) in chemotype III. All EO chemotypes of *L. graveolens* demonstrated the ability to inhibit the *A. hydrophila* growth. Thymol chemotype exhibited the strongest inhibitory effect against bacterial growth, with a minimum inhibitory concentration (MIC) of 92.4 µg/mL and a minimum bactericidal concentration (MBC) of 184.8 µg/mL. Furthermore, the results suggest that there is no synergistic or additive effect when combining different chemotypes of *L. graveolens*.

**Conclusions:** This the first report of *L. graveolens* chemotypes exhibiting antimicrobial activity against oxytetracycline-resistant *A. hydrophila*. The findings suggest the chemotype thymol could be a potential treatment for infections in the tilapia aquaculture in Guatemala.

## Resumen

**Objetivo:** El objetivo de este estudio fue evaluar la eficiencia antimicrobiana del aceite esencial de diversos quimiotipos de *Lippia graveolens* contra la cepa *Aeromonas hydrophila* resistente a la oxitetraciclina, la cual afecta principalmente a la acuicultura de tilapia (*Oreochromis* sp) en Guatemala.

**Metodología:** *L. graveolens* se colecto en tres departamentos de Guatemala, los aceites esenciales (AE) se obtuvieron mediante hidrodestilación y se caracterizaron mediante cromatografía de gases y espectrometría de masas (GC/MS). Posteriormente, se realizó un ensayo antimicrobiano utilizando pruebas de susceptibilidad en disco y dilución, y se evaluaron las interacciones sinérgicas entre los diferentes quimiotipos. Cada prueba se repitió tres veces.

**Resultados:** El análisis evidenció la presencia de veintisiete compuestos en los AE obtenidos de los quimiotipos, siendo los monoterpenos la clase principal. Los principales constituyentes identificados fueron cis-dihidro-β-terpineol (8.84 %) en el quimiotipo I, carvacrol (51.82 %) en el quimiotipo II y timol (79.62 %) en el quimiotipo III. Todos los AE de los diferentes quimiotipos de *L. graveolens* demostraron capacidad para inhibir el crecimiento de *A. hydrophila*. En particular, el quimiotipo timol obtuvo el efecto inhibitorio más fuerte contra el crecimiento bacteriano, con una concentración mínima inhibitoria (CMI) de 92.4 µg/mL y una concentración mínima bactericida (CMB) de 184.8 µg/mL. Los resultados sugieren que no existe un efecto sinérgico o aditivo al combinar los diferentes quimiotipos de *L. graveolens*.

**Conclusiones:** Este estudio constituye el primer reporte sobre la actividad antimicrobiana de los diferentes quimiotipos de *L. graveolens* contra *A. hydrophila* resistente a la oxitetraciclina. Los hallazgos sugieren que el quimiotipo timol podría ser un tratamiento potencial para las infecciones en la acuicultura de tilapia en Guatemala.

## Introduction

In aquaculture practices in Guatemala, antibiotics have primarily been utilized for therapeutic and prophylactic control of bacterial diseases. Many farmers believe that administering antibiotic therapy can provide significant advantages, such as reducing mortality rates resulting from bacterial infections. However, the repeated utilization of antibiotics has resulted in the emergence of drug-resistant bacteria, such as *Aeromonas hydrophila* (García-Pérez et al., 2021). This bacterium can bioaccumulate in fishery products through the food chain and be consumed by the local community, causing bacteria resistance or harm to human health (Rigos et al., 2004; Romero et al., 2012; Alanazi et al., 2021). As a result, to manage drug-resistant bacteria, it is necessary to either increase drug dosages or develop a new generation of antibiotics. To prevent this serious issue, the livestock must change the practice to treat diseases in fish or new alternative has emerged as an additional tool to control both resistant and non-resistant pathogens, as a means to reduce antibiotic usage and medicinal plants can be one such alternative (Olusola et al., 2013; Ham et al., 2020).

The genus *Lippia* (Verbenaceae) is among the most used plants in Central and South America for this purpose (Pascual et al., 2001). This genus comprises around 200 species of herbs, shrubs, and small trees (Terblanché and Kornelius 1996). In Guatemala, *Lippia* is represented by around 13 species, with *L. graveolens, L. alba, L. salamensis, L. chiapasensis, L. dulcis*, and *L. cardiostegia* being the most widely distributed (Standley & Williams, 1970). Ethnobotanically, *L. graveolens* is commonly referred to as Mexican oregano. It is an aromatic perennial shrub that can grow up to 2 meters tall, with short-pilose branches and leaves on petioles typically measuring 5-10 mm in length (Standley & Williams, 1970).

The shrub is characterized by its axillary capitate spikes inflorescences, which consist of white, sessile and zygomorphic flowers. These flowers are hermaphrodite and self-compatible (Ocampo-Velázquez et al., 2009). Essential oils are found in all parts of most *Lippia* genus plants, but leaves or aerial parts are particularly rich in this component, as indicated by Pascual et al. (2001). *Lippia graveolens* is typically harvested from wild populations growing in a range of ecological conditions, from semi-arid to sub-humid lands. This can result in a great morphological variability and chemical polymorphism, with essential oil composition being the most affected component (Tezara et al., 2014).

In Guatemala, researchers have identified different chemotypes of essential oils, namely carvacrol, thymol, and a minor type known as sesquiterpene (E)-caryophyllene. The latter is sometimes referred to as a mixed chemotype because it contains a variable combination of metabolites (Pérez-Sabino et al., 2012; Salgueiro et al., 2003). However, it is unknown whether the other chemotype with sesquiterpene as the major compound exists in Guatemala. To date, no research has been conducted on the antimicrobial properties of the EO chemotypes originated from wild populations of *L. graveolens* in Guatemala.

However, *L. graveolens* from various regions around the world has been investigated as a potential source of bioactive compounds possessing antibacterial and antioxidant properties, as well as for its potential use in preventing fish diseases (Bautista-Hernández et al., 2021; García-Pérez et al., 2019; Leyva-López et al., 2017). Several studies have evaluated the inhibitory effects of *L. graveolens* against a variety of bacteria, including Gram-positive strains such as *Streptococcus agalactiae, Staphylococcus aureus, S. epidermidis, Sarcina lutea, Bacillus subtilis, Enterococcus faecalis* as well as Gram-negative bacteria like *Aeromonas hydrophila, A. sobria, Vibrio cholerae, Escherichia coli, Pantoea agglomerans, Enterobacter aerogenes* (Arana-Sánchez et al., 2010; Castellanos-Hernández et al., 2020; Hernández et al., 2009; Martínez et al., 2021). Therefore, the aim of this study was to assess the antimicrobial properties of diverse chemotypes of *L. graveolens* grown wild in Guatemala, both individually and in combination, against the predominant bacterium in finfish aquaculture in Guatemala.

## Methodology

### Plant collection

Samples of a wild population of *Lippia graveolens* (Verbenaceae) in Guatemala were obtained: I. El Subinal, Guastatoya, El Progreso (N14°51LJ15 lJ / W090°08LJ05 lJ [440 m.a.s.l]), II. El Carrizal, San Jacinto, Chiquimula (N14° 37LJ18 lJ / W089° 28LJ58 lJ [700 m.a.s.l]) and III. Casas de Pinto, Río Hondo, Zacapa (N15°01LJ25 lJ / W089°36LJ35 lJ [180 m.a.s.l]). Its taxonomic name was verified using the Flora of Guatemala Part IX (Standley & Williams, 1970).

The samples were authenticated by Mr. Max Merida from San Carlos University, voucher specimens (USCG 48320, USCG 48321 and USCG 48322) were deposited at the Herbarium USCG CECON-USAC. The plant material was oven-dried at 45 LJC for 48 h, then ground into powder with an electric grinder. Then, the powder was sieved through a 300-500 µm mesh size and stored in a sealed bag at room temperature until further analysis.

### Essential oils extraction and Gas chromatography–mass spectrometry (GC/MS) analysis to determine the chemotype of *L. graveolens*

A Clevenger type device was used to extract essential oils (EO) by hydro-distillation (Majolo et al., 2018). For each extraction, 100 g of plant-powder was added in a 2,000 mL volumetric flask and covered with tap water until submerged. The extraction lasted 2 h after water boiling and the collected EO were stored at 4 LJC in 10 mL amber glass bottles. This procedure was repeated three times. The EO yield was estimated as Y (%) = M/MV x 100, where M is the mass of EO obtained (g), and MV is the amount of dried plant-powder used (g). The GC/MS analyses were performed using a Shimadzu 2010 Plus system coupled with a Shimadzu QP-2010 Plus selective detector (MSD), equipped with a DB5-MS capillary fused silica column (30 m, 0.25 mm I.D., 0.25 µm film thickness). The oven temperature was set to increase from 60 °C to 246 °C at 3 °C/min and then remained at 246 °C for 20 min. He (99.999%) was used as carrier gas with a flow rate of 1.03 mL/min, and a split ratio of 1:50. Mass spectra were taken at 70 eV. The m/z values were recorded in the range of m/z 40–700 Da. The EO components were identified by comparing their mass spectra and retention indices to literature values (Adams, 2007). Relative amounts of components were calculated based on GC peak areas without correction factors.

### Antimicrobial assay

#### Bacterium strain and culture

*Aeromonas hydrophila* (strain: CEMA150219) was obtained from the bacterial collection of the Centro de Estudios del Mar y Acuicultura, Universidad de San Carlos de Guatemala. This strain of bacterium was previously found in sick tilapia (*Oreochromis* sp.) and has shown resistance to oxytetracycline, which is the most widely used antibiotic in tilapia farming in Guatemala. To activate the strain, it was according to the National Committee for Clinical Laboratory Standards (NCCLS) standard method M31-T and considering the changes made by Alderman & Smith (2001). The bacterium was seeded on TSA agar (TSA: Merck®) for 18 hours at 28°C. The temperature was selected according to the criteria of Noga (2010), which states that for the microbiology of fish pathogens, it should be carried out at the temperature at which the organisms are naturally found, rather than 37 °C, as indicated by the standard methodology. This is because tropical or subtropical fish pathogens can exhibit poor growth or fail to grow under such conditions. Subsequently, 3-5 colonies were identified and placed in 1X phosphate-buffered saline (PBS) until reaching a bacterial concentration of 1-2 x 10^8^ CFU/mL, which was verified with an absorbance of 1.00 at a wavelength of 600 nm, measured in a spectrophotometer.

#### Disk diffusion method

The effect of different EO chemotypes: a) cis-Dihydro-β-terpineol [I], b) carvacrol [II], c) thymol [III]) and their combination of two EO chemotypes: d) [I : II], e) [I: III], f) [II:III], in a 50:50 ratio (v / v) and combining the three EO chemotypes g) [I:II:III] in a ratio of 33:33:33 (v/v)) from *L. graveolens* on *A. hydrophila* was tested using the disk diffusion method according to Bauer et al., (1966) and considering the changes made by Mazumder et al., (2020). Within 15 minutes of preparation of the inoculum, a 100 µL aliquot of *A. hydrophila* strain was evenly spread on separate Muller-Hinton agar plates (MHA: Merck®). The aliquot was spread across the agar plate using an L-shaped loop, ensuring that it covered the entire diameter of the Petri dish. Next, in an empty paper disc (BBL: Sensi-Disc®) with a diameter of 6 mm were inoculated with a total of 10 µL of each chemotype alone, in combination and oxytetracycline discs (40 µg) were used as a positive control. Each EO and combination was tested in triplicate and placed one disc per Petri dish. The plates with discs were incubated at 28 LJC for 24 hours under aerobic conditions, and after the inhibition zone diameters (IZD) were measured with electronic calipers with an accuracy of 0.01 mm. The interpretation of the antibacterial activity of EO was determined according to the criteria established by García-Pérez & Marroquín-Mora (2020), considering a specific plant extract, essential oil, or other natural product is considered to have antibacterial action when it produces IZD equal to or greater than 50% compared to the most used antibiotic. In the case of Guatemala, the most used antibiotic in aquaculture is oxytetracycline.

#### Minimum inhibitory (MIC) and minimum bactericidal concentrations (MBC)

MIC and MBC determinations were done using the broth macrodilution assay according to NCCLS M31-T, considering the changes made by Alderman & Smith (2001). Two-fold serial dilutions of each essential oil chemotype of *L. graveolens* were prepared in the Muller-Hinton broth (MHB: Merck®) with a concentration ranging from 11.5 – 11,825 μg/mL. Triton X (Sigma-Aldrich®) was used as the EO solubilizer at a concentration of 0.1% (v/v). The *A. hydrophila* inoculum was adjusted to a concentration of 1 x 10^8^ CFU/mL in PBS and then transferred into MHB to obtain a bacterial count of 1 x 10^5^ CFU/mL. The tubes were then incubated at 28 LJC for 24 hours under aerobic conditions. Finally, an aliquot of 100 µL from each tube was plated in the TSA and incubated for 18 hours for bacterial count. The MIC and CMB were established according to Levison (2004) criteria: MIC was defined as the minimal concentration of antibiotic that prevents the clear suspension of 10^5^ CFU/mL from becoming turbid after overnight incubation; turbidity generally connotes a 10-fold increase in bacterial density. And MBC was determined as the lowest concentration of antibacterial agent that reduces the viability of the initial bacterial inoculum to ≥ 99%.

#### Assessing synergistic interaction among chemotype of L. graveolens

The synergistic interaction among *L. graveolens* EO chemotypes was determined according to Nikkhah et al., (2017) and Gutierrez et al., (2008) with modifications. Four interactions were tested by combining the three EO chemotypes: a) I:II, b) I:III, c) II:III, in a ratio of 50:50 (v/v) and combining the three EO chemotypes d) I:II:III in a ratio of 33:33:33 (v/v). Once the independent MIC was determined, the fractional inhibitory concentration (FIC) was calculated as follows:

- FIC of chemotype cis-Dihydro-β-terpineol (FIC_I) = MIC (cis-Dihydro-β-terpineol_[a, b]_) in combination/MIC (cis-Dihydro-β-terpineol) alone.
- FIC of chemotype carvacrol (FIC_II) = MIC (carvacrol_[a,c]_) in combination/MIC (carvacrol) alone.
- FIC of chemotype thymol (FIC_III) = MIC (thymol_[b,c]_) in combination/MIC (thymol) alone.
- FIC of all chemotypes (FIC_IV) = MIC (all chemotypes [d]) in combination/MIC (each chemotype) alone.

The FIC Index (FICi) was calculated as the sum of each FIC. The results obtained were interpreted as follows: synergistic effect (FICi LJ≤LJ0.5); additive effect (0.5< FICi ≤1); no interactive effect (1< FICi ≤4); antagonistic effect (FICIi >LJ4) as described by Gutierrez et al., (2008).

### Data analysis

The EO yields and inhibition zone diameters for disk diffusion assay were statistically analyzed with a nonparametric test of Kruskal Wallis followed by Mann– Whitney–Wilcoxon test (*p* < 0.05). The major compounds of the EO chemotypes from *L. graveolens* identified (greater than 1%) were used to determine the chemotaxonomy by using a Principal Component Analysis (PCA). Statistical analysis was carried out using statistical software (R Core Team, 2022).

### Analysis and results

The EO extracted from *Lippia graveolens* collected in Guatemala showed a yellow-reddish color, and yields were significantly different among departments (*p* < 0.05): 1.42 ± 0.25 % for El Progreso, 1.64 ± 0.14% for Chiquimula and 3.77 ± 0.28 % for Zacapa. Therefore, it may be inferred that there is a relationship between EO origin and its total production. These findings coincided with Martínez-Natarén et al. (2014), who found that the EO yield of different plants varied according to the edaphoclimatic gradient. Therefore, different odors, colors, and amounts of metabolites may be found. Concerning the chemotype aroma, there was an evident difference, with the EO chemotypes from Zacapa and Chiquimula having a spicy and an intense pungent odor, while El Progreso chemotype had a less intense pungent odor. Accordingly, based on odor, color, and consistency, it is inferred that the number of phenolic compounds may be greater in the Zacapa and Chiquimula chemotypes than in El Progreso. On the contrary, El Progreso could have a greater amount of non-phenolic substances (oxygenated sesquiterpenes). The chromatographic profiles of each chemotype of *L. graveolens* are shown in Figure 1.

**Figure 1.**
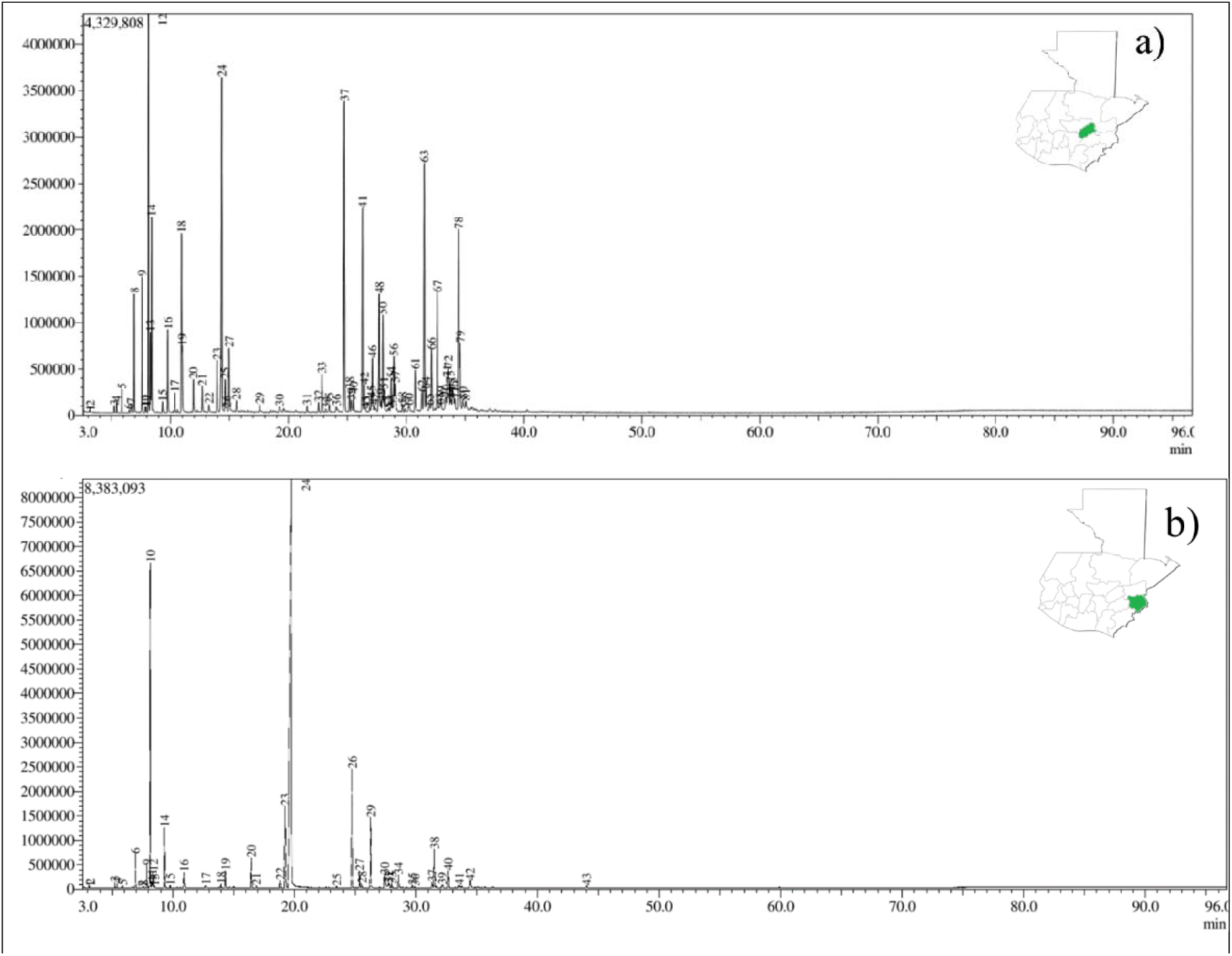
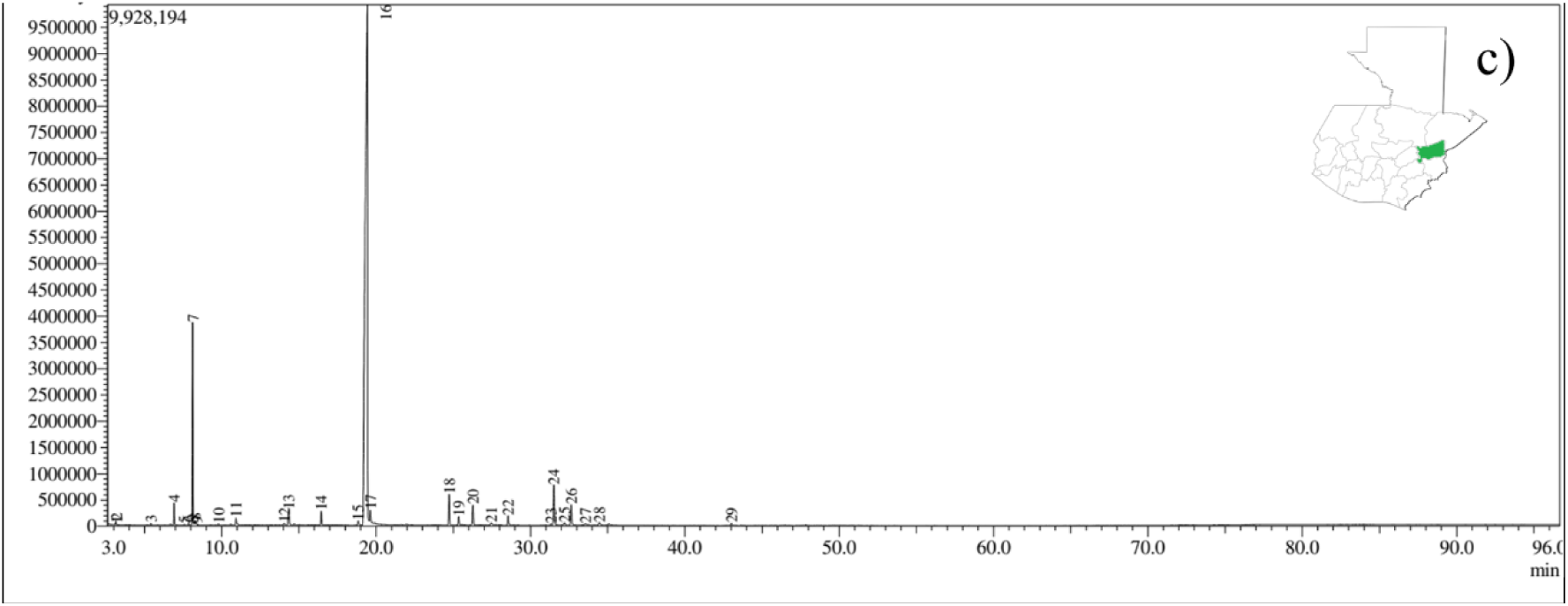
Derived from research. GC chromatogram of essential oils from *Lippia graveolens* collected from: a) El Progreso (I: *cis*-Dihydro-β-terpineol), b) Chiquimula (II: Carvacrol) and c) Zacapa (III: Thymol) departments.

According to table 1, the content and proportion of chemical constituents of the EO (monoterpenes, sesquiterpenes, phenylpropanoids and other compounds) were different among departments. The PCA horizontal axis explained 81.4% of the total variance and the vertical axis 18.6% (Figure 2), suggesting the existence of three chemotypes (I: *cis*-Dihydro-β-terpineol: El Progreso, II. Carvacrol: Chiquimula, and III. Thymol: Zacapa) characterized by their different content of chemical metabolites.

**Figure 2.**
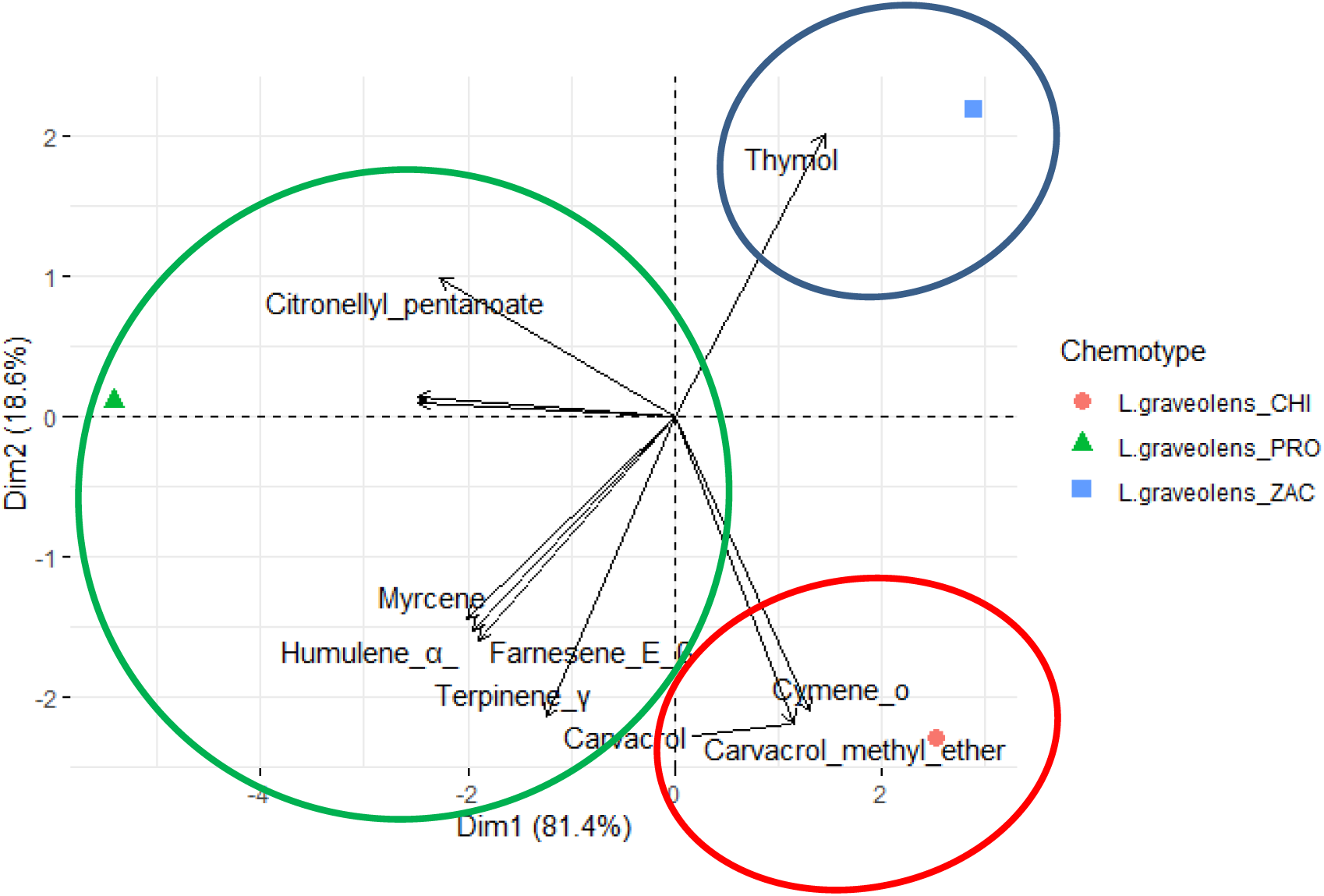
Derived from research. PCA of the 15 major components of the chemotypes among the departments PRO: El Progreso, CHI: Chiquimula and ZAC: Zacapa, of *Lippia graveolens*.

**Table 1.**
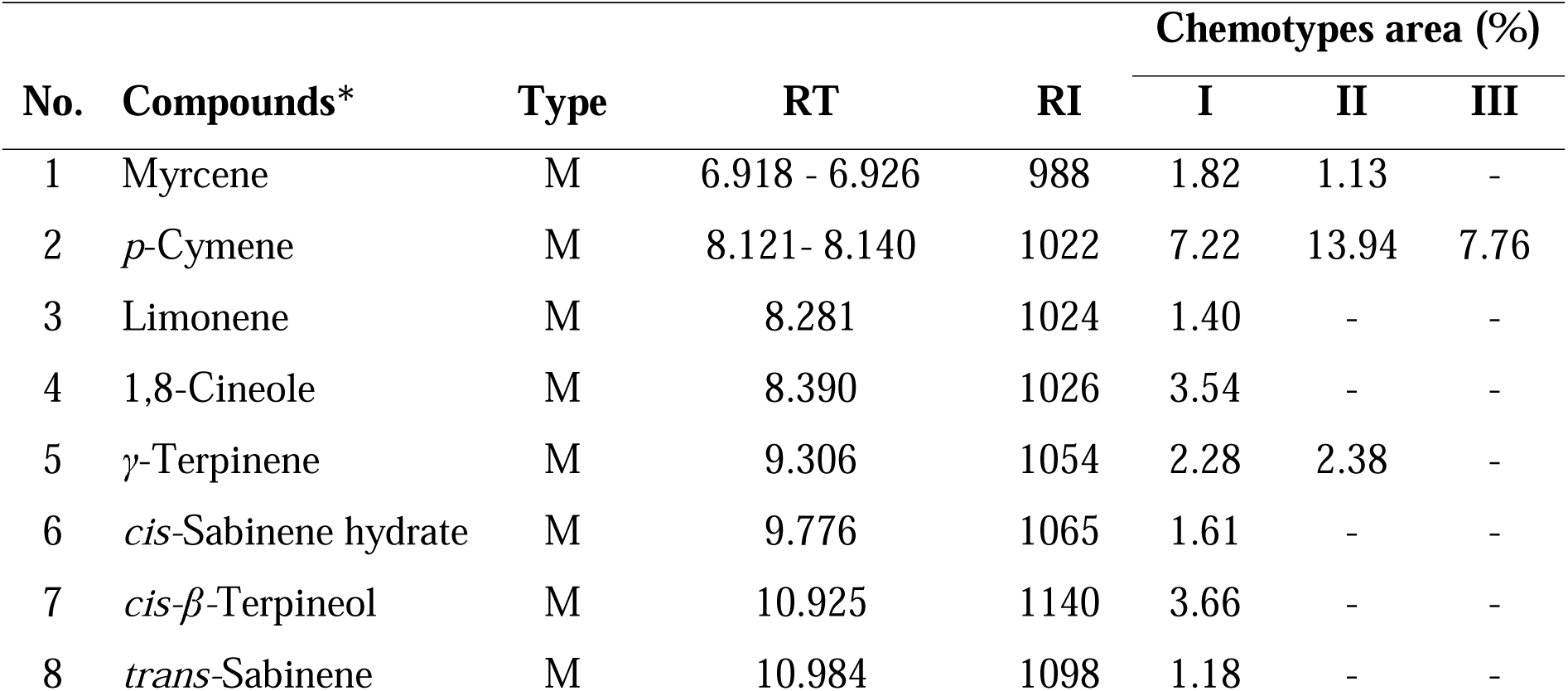
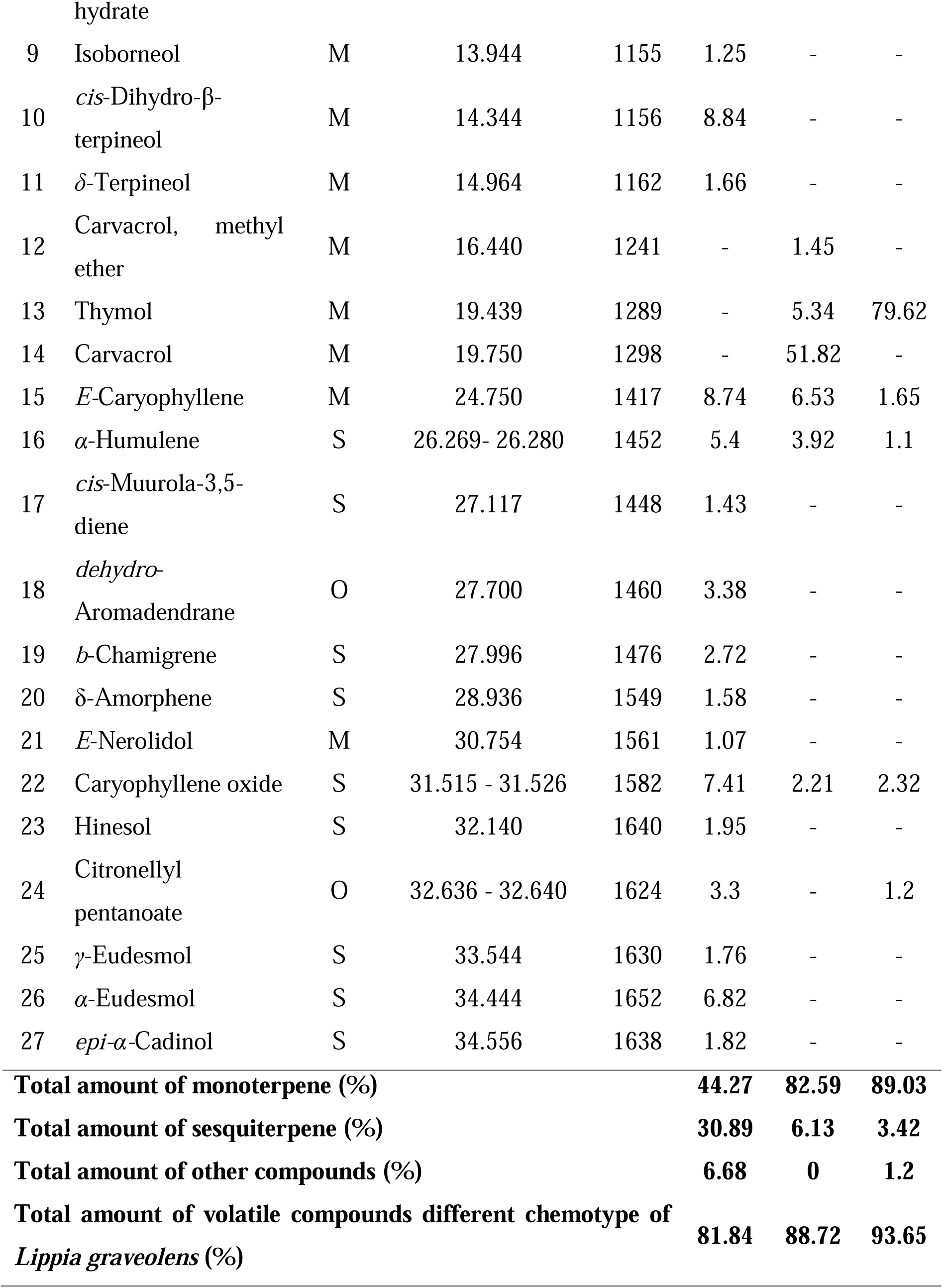
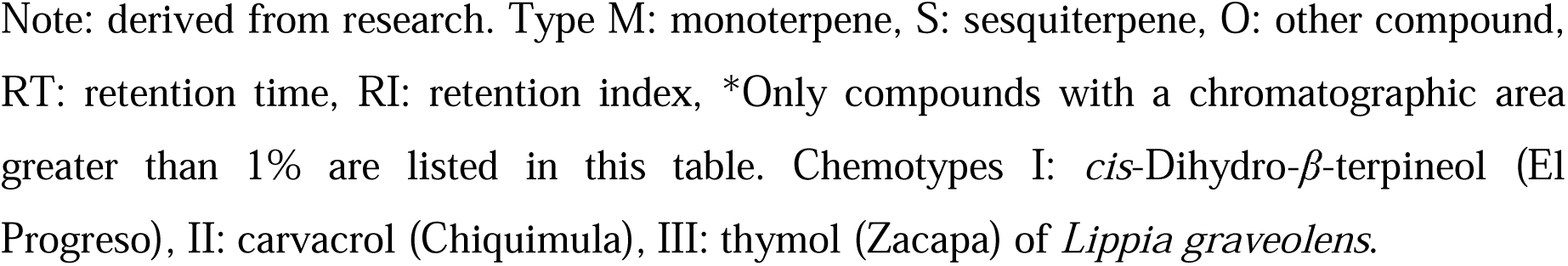
Chemical composition (%) of the three chemotype essential oils from Lippia graveolens.

In general, twenty-seven compounds were identified in the EO of *L. graveolens* (Table 1), which accounted for 81.84%, 88.72%, and 93.65% of the total EO composition of chemotypes I, II, and III, respectively. The monoterpenes constituted the main class with *cis*-Dihydro-β-terpineol (8.84%) as a major constituent of chemotype I, carvacrol (51.82%) and thymol (79.62%) as the major constituents of chemotype II and III, respectively. The second major group of compounds found in the EO were the sesquiterpenes, with caryophyllene oxide as the major constituent of chemotypes I (7.41%) and III (2.32%), whereas the α-Humulene (3.92%) was the major constituent of chemotype II. These results evidenced the EO chemical polymorphism of this species as mentioned previously for Guatemala (GT) and México (MX): thymol (GT: 67.4 – 73.5% / MX: 84.76 - 91.77%), carvacrol (GT: 50.3-51.8% / MX: 72.20 - 83.49%), (E)-caryophyllene (GT:10.3 – 12.3 % / MX: 0%), and caryophyllene oxide (GT: 2.7 – 4.9% / MX: 4.94 - 11.34%) (Martínez-Natarén et al., 2011; Pérez-Sabino et al., 2012; Salgueiro et al., 2003; Senatore & Rigano, 2001). Pérez-Sabino et al., (2012) classified the chemotype I, as mixed because its heterogeneous compounds summed no more than 15% of the chemical structure. In this study, chemotype I had *cis*-Dihydro-β-terpineol (8.84%), *E-*Caryophyllene (8.74%) and *p*-Caryophyllene oxide (7.41%) as major constituents of the EO. These proportions are different from those reported previously from EO obtained in the same location. The differences in terms of some compounds could be attributed to genetic plasticity of *L. graveolens*, soil composition, climate variability, harvest time and extraction method among other factors (Benjemaa et al., 2022; Martínez-Natarén et al., 2014).

The results indicate that three EO chemotypes of *L. graveolens* provide promising alternatives for controlling the growth of *Aeromonas hydrophila* isolated from tilapia. Chemotypes III and II were the most effective, presenting IZD ranging from 28.46 to 34.03 mm (Table 2). Chemotype I exhibited the smallest IZD (9.95 mm), although it is important to note that even though the IZD is the lowest, it still indicates an effect based on the criteria established in this study. For the combination of EO chemotypes, the highest IZD (29.19 – 33.43 mm) was for II:III followed by I:III and I:II:III. The lowest IZD (24.82 mm) was obtained for the combination of chemotype I:II. The positive control (OXI) had an IZD of 10.32 mm, indicating the resistance of the *A*. *hydrophila* to oxytetracycline.

**Table 2.**
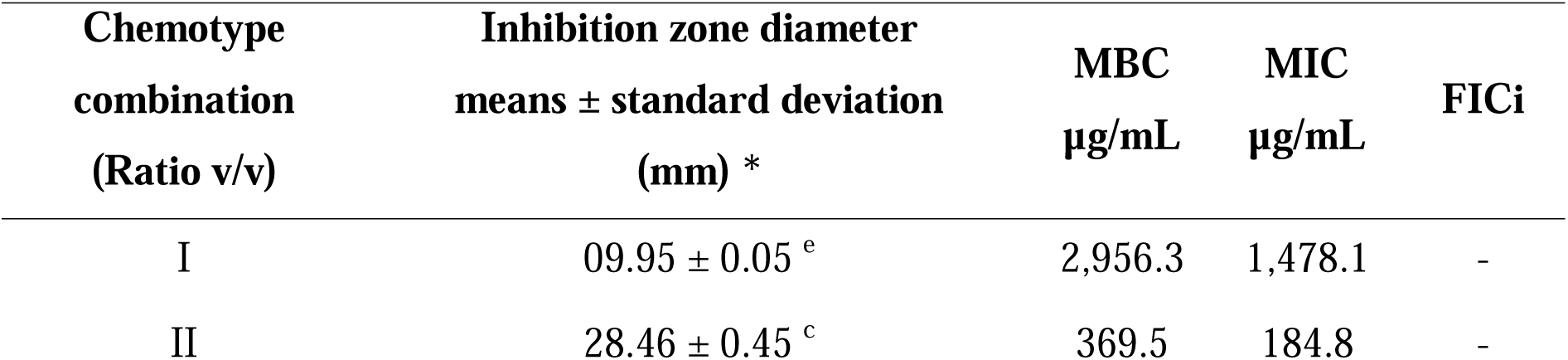
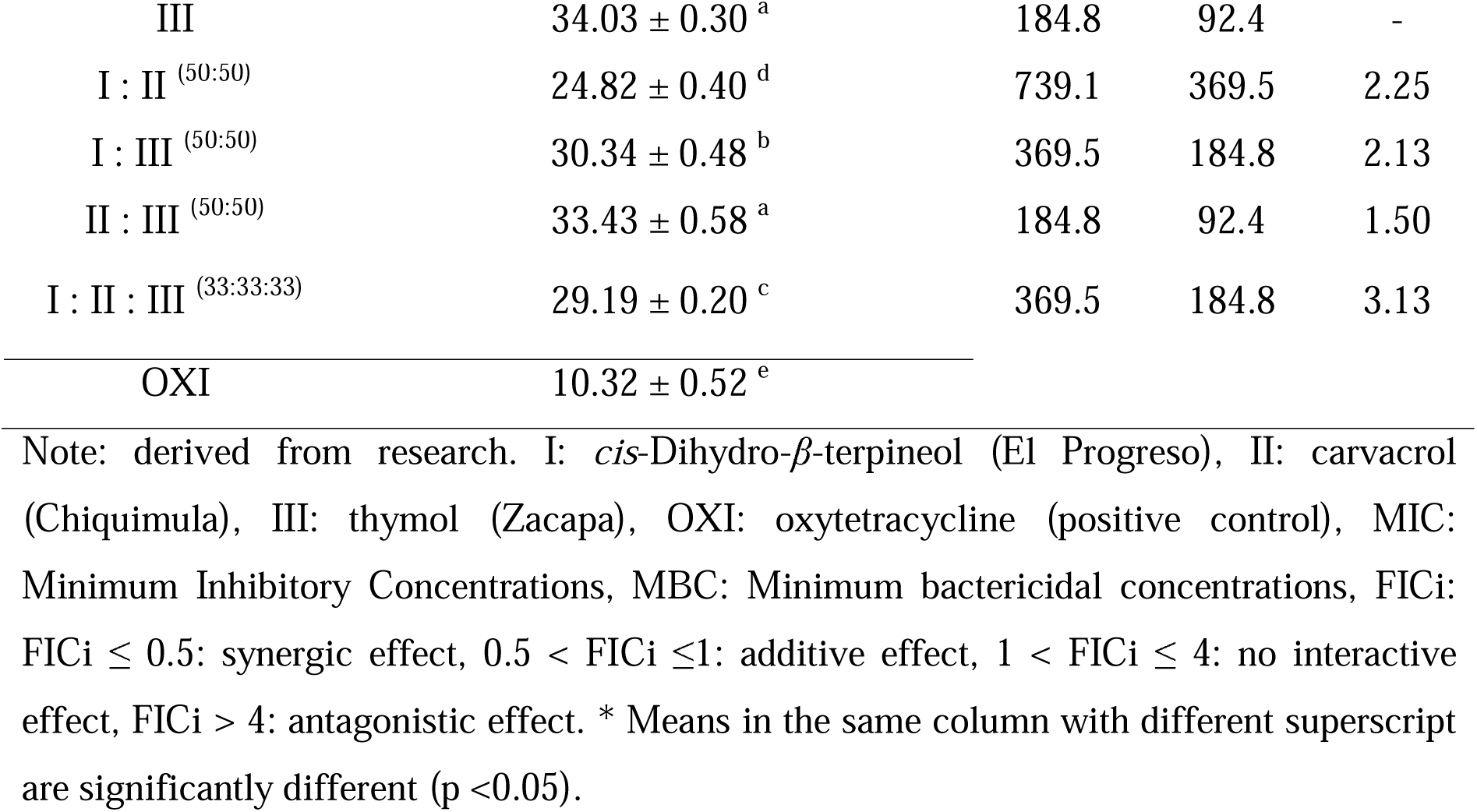
Antibacterial activity of the three chemotype essential oils from *L. graveolens* against *A. hydrophila*.

The MIC and MBC values for the three EO chemotypes ranged from 92.4 – 2,956.3 µg/mL (Table 2). The lowest MIC and MBC values found were for chemotype III, 92.4 – 184.8 µg/mL, followed by chemotype II: 184.8 – 369.5 µg/mL and finally chemotype I: 1,478.1 – 2,956.3 µg/mL. Although chemotype I obtained the highest MIC and MBC, it still may be considered effective; according to Bussmann et al., (2010), MIC values below 5,000 µg/mL are considered to exert strong antimicrobial activity.

Results indicated that the three EO chemotypes mixture from *L. graveolens* did not show a synergistic or additive effect. However, the greatest effect was observed when chemotype III was combined with any of the other chemotypes (I and II), even with chemotype I (with the lower inhibition capacity), thus having an antibacterial effect. The EO extracted from chemotype III (thymol) exhibited better antibacterial properties than EO components of chemotypes II and I, with a significantly greater inhibition zone diameters and a smaller concentration for MBC and MIC. Therefore, it may be inferred that the properties of thymol metabolite could be responsible for the antibacterial activity observed in this study. It is well documented that *L. graveolens* have an antibacterial effect against a broad group of pathogens, especially against Gram-negative bacteria (Arana-Sánchez et al., 2010; Bautista-Hernández et al., 2021; Hernández et al., 2009; Leyva-López et al., 2017; Salgueiro et al., 2003). However, this is the first report analyzing the effect of different EO chemotypes from *L. graveolens* growing in Guatemala against aquaculture oxytetracycline resistant bacteria, such as *A*. *hydrophila*. It showed that secondary metabolites of *L. graveolens* may have a direct effect as antimicrobial agents, as stated by several authors for other plant species (Höferl et al., 2020; Koba et al., 2009; Nwanosike et al., 2016).

According to the results, most of the antimicrobial effects of the EO tested may be attributed to the presence and proportions of thymol and carvacrol in the three chemotypes, both acting independently, and probably having the same action pathway, since there was no proved synergic or additive effect.

Although the antimicrobial effects of *L. graveolens* is well established, the mechanism of action is mostly unknown. Based on what was observed in the study of Lambert et al. (2001), it is inferred that the essential oil of *L. graveolens* can rupture the external membrane and may cause permeability of the ATP; this derived from the fact that the main components for *L. graveolens* is thymol or carvacrol. Furthermore, it was found that thymol and carvacrol could disintegrate the outer membrane of bacteria of *A. hydrophila*, isolated from different sources, and other types of bacteria, such as *Escherichia coli* and *Salmonella typhimurium* (Helander et al., 1998). On the other hand, it is deduced that the components of chemotype I showed lower antibacterial activity due to its relative amount and type of sesquiterpenes, which have weak antimicrobial activity. This weak antimicrobial activity has been attributed to the fact that most sesquiterpenes does not have the appropriate hydrophobicity to break down the bacterial membrane (Bassole et al., 2005).

## Conclusions

The results suggest the essential oil from *L. graveolens* offer a promising alternative for the control of *Aeromonas hydrophila* growth. It was found three chemotypes: I: cis-Dihydro-β-terpineol located in El Progreso Department, II: Carvacrol in the Chiquimula Department, and III: Thymol in the Zacapa Department. All EO exhibit antibacterial activities even in different proportion, but there is no synergistic or additive effect when combining different chemotypes. The results suggest that the EO chemotype thymol can be used for aquaculture bacterial infections. However, further studies are needed to assess the toxicity of *L. graveolens* EO on tilapia. Because certain EO chemicals may have a direct impact on the target bacteria, larger amounts could potentially become toxic to fish and other organisms. Furthermore, it is recommended to investigate the *in vivo* efficacy of EO treatment for *Aeromonas* infections in finfish production in Guatemala.

## Acknowledgements

This research constitutes a chapter of the first author’s doctoral thesis of the program: ‘‘Doctorado en Ciencias Naturales para el Desarrollo (DOCINADE), Instituto Tecnológico de Costa Rica’’, and it was funded by the Universidad de San Carlos de Guatemala cohort 2017. I would like to express my very great appreciation to Carolina Marroquín for the accurate review of this document.

## Conflict of Interest

The authors declare no competing interests.

## Author contribution statement

All the authors declare that the final version of this paper was read and approved. The total contribution percentage for the conceptualization, preparation, and correction of this paper was as follows: J.R.G.P 50%, J.F.P.S 10%, S.E.M.E 10% A.R.S. 10% and J.B.U.R. 20%.

## Data availability statement

Please select only one of the following: The data supporting the results of this study will be made available by the corresponding author, **[J.R.G.P]**, upon reasonable request.

## Notes

### Competing Interest Statement

The authors have declared no competing interest.

